# Capturing the complexity of topologically associating domains through multi-feature optimization

**DOI:** 10.1101/2021.01.04.425264

**Authors:** Natalie Sauerwald, Carl Kingsford

## Abstract

The three-dimensional structure of human chromosomes is tied to gene regulation and replication timing, but there is still a lack of consensus on the computational and biological definitions for chromosomal substructures such as topologically associating domains (TADs). TADs are described and identified by various computational properties leading to different TAD sets with varying compatibility with biological properties such as boundary occupancy of structural proteins. We unify many of these computational and biological targets into one algorithmic framework that jointly maximizes several computational TAD definitions and optimizes TAD selection for a quantifiable biological property. Using this framework, we explore the variability of TAD sets optimized for six different desirable properties of TAD sets: high occupancy of CTCF, RAD21, and H3K36me3 at boundaries, reproducibility between replicates, high intra- vs inter-TAD difference in contact frequencies, and many CTCF binding sites at boundaries. The compatibility of these biological targets varies by cell type, and our results suggest that these properties are better reflected as subpopulations or families of TADs rather than a singular TAD set fitting all TAD definitions and properties. We explore the properties that produce similar TAD sets (reproducibility and inter- vs intra-TAD difference, for example) and those that lead to very different TADs (such as CTCF binding sites and inter- vs intra-TAD contact frequency difference).

## 1 Introduction

Since the introduction of a sequencing technique to study the genome-wide three-dimensional structure of chromosomes [15], many studies have shown a connection between this architecture and regulatory mechanisms. Genome structure plays a role in gene regulation [14], cell cycle coordination [7], and many human diseases and disorders (see [25] for review). This structure has been partially described by topologically associating domains (TADs), the building blocks of genome architecture that are regions of the chromosome interacting more highly within themselves than with other regions [8]. TADs in particular have been suggested to bring together target genes with their intended regulatory elements [9] and insulate them from interactions with other nearby enhancers and promoters, though their exact role and even definition remains an area of active study [2].

One of the main challenges in describing and identifying TADs is a lack of consensus on how to define these structures, with different features used to define TADs by different analyses. Computationally, we expect TADs to have dense intra-TAD contacts, sparse inter-TAD contacts, and show a strong shift in contact direction at TAD boundaries, among other features [2, 8]. TADs are generally identified through optimization of one of these computational properties. One TAD finder, Armatus [10], finds the TAD set with the greatest sum of TAD densities, where TAD density is a scaled sum of contact counts within each TAD. TopDom [24] and Insulation Score [4], both successful TAD finders, instead set TAD boundaries at the minima of a function summing the contacts within a window centered on each genomic section, thereby identifying TADs as the regions in between highly insulated sections. The first computational method for TAD finding, called Domain Caller [8], was based on quantifying the direction of contact bias, or whether a chromosome segment preferentially interacted with other segments upstream of it or downstream, and used a Hidden Markov Model to define TADs where this directionality index switched from upstream to downstream. These concepts are all related and have all been successful in TAD identification, though no one method performs best across all evaluation metrics [11, 5, 29].

Without a gold standard or ground truth TAD set to which we can compare computational predictions, we assess TAD sets based on various properties that we expect from them. Biologically, TADs should show increased binding of CTCF, certain histone markers, and cohesin at their boundaries, and remain fairly consistent across replicates [8, 10, 20]. CTCF is a critical structural protein for TAD boundaries, and we therefore expect to see an enrichment of CTCF binding at these boundary locations [8]. Similarly, the histone marker H3K36me3 has been shown to be enriched at TAD boundaries, to the extent that it can be combined with other histone markers to predict TAD locations [10, 22]. Because of the significant involvement of cohesin in TAD formation, two protein components of cohesin, RAD21 and SMC3, have also been shown to peak around TAD boundaries [24]. While TADs have been shown to vary between single cells [28], on the population level they are expected to remain fairly consistent between replicates [20]. On a fundamental level, TADs are defined by increased contact frequency within their interiors and somewhat depleted contact frequency between different TADs. This can be quantified by looking at the distributions of inter-TAD contacts and intra-TAD contacts, with the expectation that the distribution of intra-TAD contacts is generally higher than that of inter-TAD contacts.

Several reviews have assessed the performance of TAD finders on their ability to fit these biological properties, always finding that no one method captures every property equally well, and there is a significant amount of disagreement in the TADs output by the different methods [11, 5, 29]. It is unclear why these different computational TAD definitions lead to such different TAD sets, and why some perform better on certain metrics than others. For example, while Armatus TADs are fairly reproducible [11] and show higher intra-TAD contacts than inter-TAD contacts [5], the boundaries do not contain as many CTCF, RAD21, or SMC3 peaks as other tools [29]. On the other hand, Domain Caller TADs have very different inter-TAD and intra-TAD contact distributions [5], align extremely well with CTCF, RAD21, and SMC3, but do not have particularly impressive histone marker measures [29] and have very low reproducibility [11]. TopDom and Insulation Score seem to occupy a space of doing fairly well across all metrics and generally agreeing well with other TAD finding results, but rarely coming out as the best TAD finder in any particular measure [11, 5, 29].

The relationship between the computational definitions that each TAD-finding method implements and the resulting TAD sets remains unclear, and it is still an open question whether there exists an algorithm to identify TADs that would be superlative across assessment metrics, or whether there are inherent tradeoffs between these evaluation criteria. To study these questions and the variability of computational TAD predictions, we have developed a general TAD finder to reflect the space of potential TAD sets that can be optimized for any quantifiable TAD property. This allows exploration of the computational definitions of TADs, and the ability to combine computational components rather than relying on a single TAD definition.

Parameters of the model can be chosen to guide TAD selection towards any quantifiable desired property, provided the relevant data. This connects the computational definitions with the biological properties we use to assess TAD sets, identifying TADs that fit a combination of computational definitions and optimize a specific biological property such as high CTCF occupancy at boundaries. We choose parameter sets to optimize six different desirable TAD properties: (1) many CTCF binding sites at boundaries, high occupancy (measured by ChIP-seq data) of (2) CTCF, (3) RAD21, or (4) H3K36me3 at TAD boundaries, (5) reproducibility, and (6) large difference in inter- and intra-TAD contact frequency. We compare the resulting TAD sets across 12 different cell and tissue types to quantify variability and study the computational and biological properties that lead to similar TAD patterns. We find that some cell types show extremely high variability in TAD sets, while others are much more consistent, and analyze which properties lead to each of these outcomes.

## 2 Methods

To reflect the space of reasonable computationally-defined TAD sets, we developed a general algorithm for TAD identification. This algorithm, called FrankenTAD, optimizes three computational TAD properties that have been used in successful TAD finders. Tuning the parameters of this algorithm can lead to very different TAD sets, so we can use various desired properties to select parameters that will guide the resulting TAD set towards the given target. Parameters are optimized for one of several possible objective functions, reflecting various biological and technical TAD properties discussed below. An overview of the method is shown in Figure 1, and all methods and analysis code can be found at https://github.com/Kingsford-Group/frankentad.

**Figure 1:**
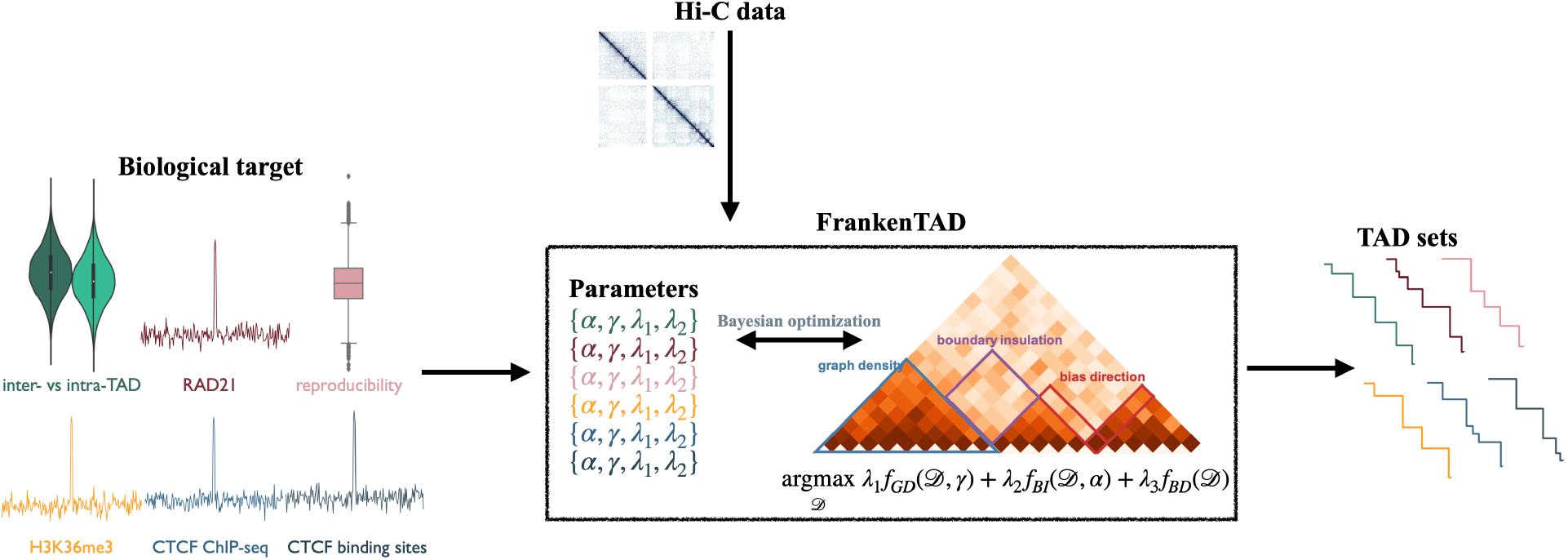
Overview of the main components of this work.

### 2.1 TAD-finding algorithm

FrankenTAD optimizes a linear combination of three TAD features: density within TADs, insulation between TADs, and a change in contact bias around TAD boundaries (see center of Figure 1 for an illustration of these properties). Varying the model parameters leads to different TAD sets that emphasize different computational and definitional aspects of TADs.

Below, 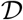 represents a set of TADs on a given chromosome, the *N* ×*N* matrix *A* is the normalized Hi-C matrix with *N* bins, each bin representing a segment of *k* bases where *k* is the data resolution, *A_i,j_* is the normalized Hi-C value between bin *i* and bin *j*, and an individual TAD can be represented by an interval [*a, b*].

The core of FrankenTAD is the following objective function, combining three different TAD definitions, where *λ* = 〈*λ*_1_, *λ*_2_, *λ*_3_〉:

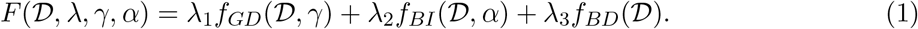

The three components of this function, *f_GD_, f_BI_,* and *f_BD_* are explained in detail below. Each component scores a different aspect of a TAD set: dense intra-TAD contacts (*f_GD_*), insulation between TADs (*f_BI_*), or a shift in bias direction at TAD boundaries (*f_BD_*).

#### 2.1.1 Graph density component

The first property included in FrankenTAD is inspired by Armatus [10], in which TADs are chosen to maximize the sum of scaled subgraph densities of each TAD. Computationally, this defines a TAD as a high density subgraph of the graph induced by the Hi-C matrix. The following objective function 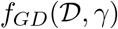 is used by FrankenTAD and Armatus:

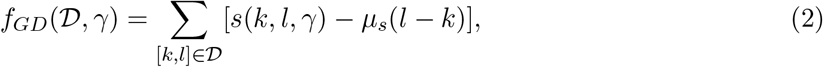

where *γ* is a parameter to be optimized, *s*(*k, l, γ*) is

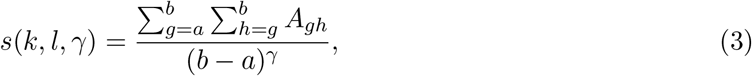

and *μ_s_*(*l* − *k*) is the mean value of *s*(*k, l, γ*) for all possible TADs of length *l* − *k*. For details on this function, see Fillipova et al. [10].

#### 2.1.2 Boundary insulation component

The next component of FrankenTAD is inspired by TopDom [24], in which TADs are defined by the strength of their boundaries rather than the density of their contacts. A sliding window is used to quantify the insulation from other nearby regions, and boundaries are identified by finding the local minima of this function. The function quantifying the average contact frequency for each bin is

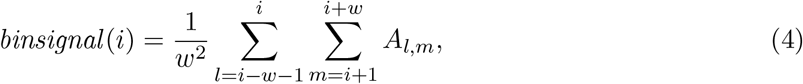

where the parameter *w* controls the window size. The expectation is for this *binsignal*(*i*) to be high if *i* is near the center of a TAD, and low if *i* is at or near a TAD boundary. Details on this function can be found in Shin et al. [24]. The following function should be maximum when TAD boundaries are at the local minima of the *binsignal*(*i*) function:

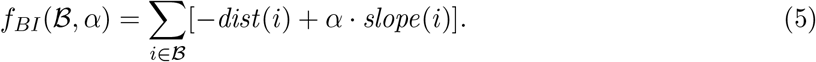

We therefore score a TAD set based on the distance from its boundaries to the nearest local minima in the *binsignal* function, and the average slope of *binsignal* around the boundary. The set 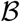 contains all TAD boundaries, *dist*(*i*) is the number of bins from *i* to the nearest local minimum, *α* is a parameter to be optimized, and *slope*(*i*) is the average slope of *binsignal* around *i*, defined as:

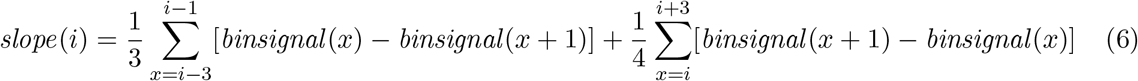

This approach should prioritize TAD boundaries not only at local minima, but those with especially high differences in insulation around the boundary.

#### 2.1.3 Bias direction component

The third component of FrankenTAD comes from the Domain Caller TAD finder [8], which is based on the insight that TADs and their boundaries will display particular contact patterns, with bins near the start of a TAD displaying a strong bias towards contacts downstream, while bins near the end of a TAD will show a strong bias towards upstream contacts. At TAD boundaries, we therefore expect to see a switch from downstream to upstream bias. This notion of contact bias was quantified in the following way in Dixon et al. [8]:

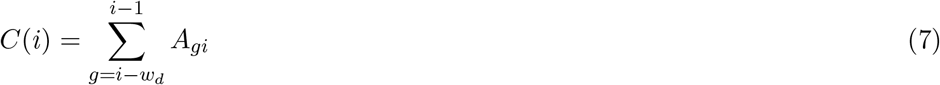

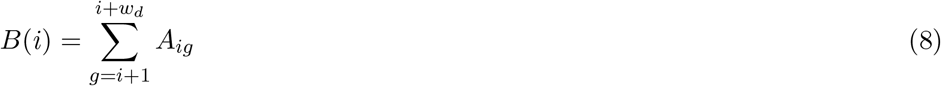

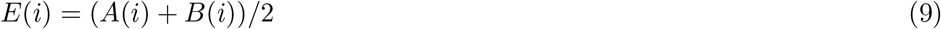

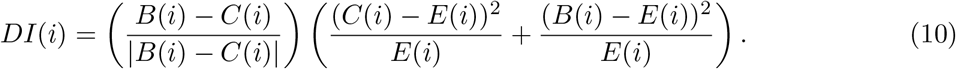

In this formula, *C*(*i*) represents the number of upstream contacts of bin *i*, *B*(*i*) represents the downstream contacts, *E*(*i*) is the average number of contacts, and *w_d_* is a window size adjusted based on the data resolution to consider contacts within 2Mb up or downstream of the bin of interest. We expect *DI*(*i*) to be strongly positive at the start of a TAD and strongly negative at the end of a TAD, leading to the following scoring function:

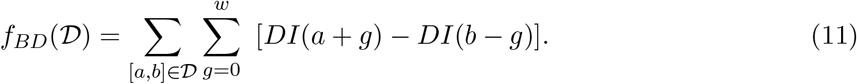

#### 2.1.4 Dynamic program to optimize multi-feature TAD definition

Ideally, TADs should fit all of these definitions: high density within a TAD, strong insulation at boundaries, a strong downstream contact bias near the start, and a strong upstream contact bias near the end of the TAD. A linear combination of these properties (Equation 1) provides a flexible framework for identifying TAD sets based on these computational features.

We identify TADs by maximizing Equation 1 through a dynamic program. For any position *b* on the chromosome, the optimal TAD set over the interval [0, *b*] is given by:

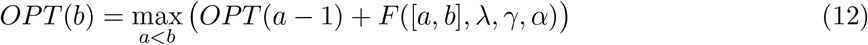

Efficiency is achieved through significant precomputation of the elements of each subfunction, as well as the above dynamic program.

### 2.2 Data-driven objective functions

There are five parameters required by FrankenTAD; one weight for each subfunction represented by *λ* = 〈*λ*_1_, *λ*_2_, *λ*_3_〉, an internal *γ* parameter for the graph density subfunction, and an internal *α* parameter for the boundary insulation subfunction. One weight can be fixed (ie, *λ*_3_ = 1), so that four parameters must be provided to run FrankenTAD. We define six different data-driven objective functions to select these parameters, thereby guiding the TAD sets we find towards these properties. Each of these properties has been used to assess the quality of TAD sets, with existing TAD finders showing clear tradeoffs between them. We formulate each property as an objective function, and parameters are chosen to find TAD sets that optimize this objective function.

#### 2.2.1 CTCF binding sites

While its exact role is unclear, the structural protein CTCF is widely understood to be critical to TAD architecture. Peaks in the number of CTCF binding sites at TAD boundaries have been used as a validation of TAD quality [10]. Binding site locations are not cell-type specific and are relatively easy to identify, making them the best way to optimize for protein locations in the absence of ChIP-seq data which may not be available for a given species, cell, or tissue type. To optimize for CTCF binding sites, we maximize a function that counts the number of binding sites at each TAD boundary in a candidate TAD set. Let *n_site_*(*i*) be the number of CTCF binding sites within bin *i*, and recall that 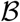 is the set of all TAD boundaries. Then

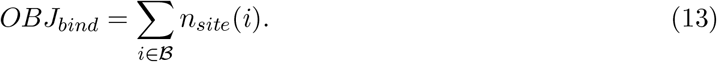

In order to find TAD sets with many CTCF binding sites at TAD boundaries, the parameters of FrankenTAD are chosen to maximize *OBJ_bind_*.

#### 2.2.2 ChIP-seq peaks: CTCF, RAD21, and H3K36me3

While binding site locations are informative of where proteins could bind, ChIP-seq data reveals the true locations of bound proteins in a specific cell or tissue type. We therefore use a similar objective function to maximize the number of peaks of bound proteins at TAD boundaries for three different structurally associated proteins. RAD21 is a protein component of the cohesin complex, which has been implicated as a key structure in TAD formation, along with CTCF. H3K36me3 is a histone marker that has also been associated with TAD structures [10, 22] and is expected to be enriched at TAD boundaries. We therefore use ChIP-seq peaks of CTCF, RAD21, and H3K36me3 to select parameters that will identify TAD sets with boundaries at the locations of binding peaks, with *n_bound_*(*i*) as a function that counts the number of ChIP-seq peaks with midpoints within the bin *i*:

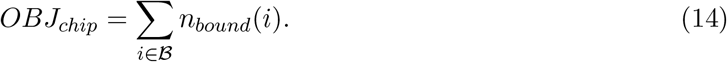

Highlighting the weakness of relying on ChIP-seq data, only 6 of 12 total cell and tissue types studied here had publicly available RAD21 ChIP-seq data, so the analysis of cohesin is more limited than the others.

#### 2.2.3 Reproducibility

There can be significant variability between single cells of the same population [26, 17, 3], but on a population level we broadly expect replicates to show similar TAD sets [20]. The Jaccard Index, 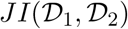, a distance metric quantifying the overlap between two sets, is used to quantify the similarity between replicate TAD sets. In this case, 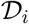 represents a TAD set, and the Jaccard Index is computed as the ratio of the intersection of the TAD sets to their union. TADs must share both starting and ending locations to be counted in the intersection of the two TAD sets. Parameters are chosen here to identify TAD sets on each replicate with maximal Jaccard Index between them:

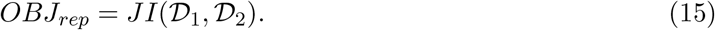

In this work we use two replicates for each cell type, but if more replicates are available this objective could be expanded as the sum of all pairs of replicate JI values.

#### 2.2.4 Inter- versus intra-TAD contact frequency

Perhaps the most fundamental property of TADs is that there are more interactions within TADs than between them, which can be quantified by comparing the inter-TAD contact frequency with the intra-TAD contact frequency. This property is the closest to the computational features optimized by FrankenTAD, compared to the biological properties reflected by ChIP-seq data. Intra-TAD contact frequency is computed as the mean of all Hi-C values within TADs in the given TAD set (Equation 16), where *n* is the total number of intra-TAD matrix entries. Inter-TAD contact frequency is computed as the mean of all Hi-C values with bins in adjacent TADs. If we describe the *t^th^* TAD along the chromosome as [*a_t_, b_t_*] (the TAD begins at bin *a_t_* and ends at bin *b_t_*), the next TAD along the chromosome can be described as [*a*_*t*+1_, *b*_*t*+1_]. The inter-TAD contact mean is given by Equation 17, where *m* is the total number of inter-TAD matrix entries considered. To identify TADs with the greatest change in these two values, we maximize their difference with *OBJ_int_*:

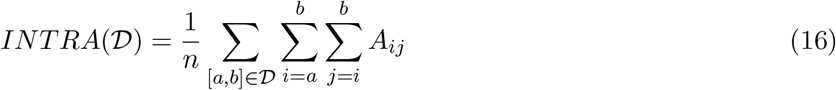

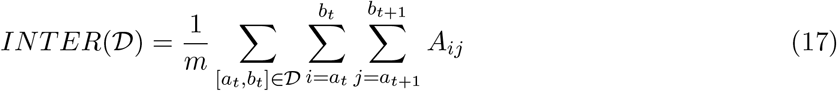

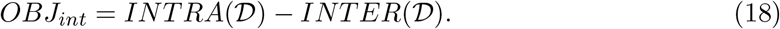

#### 2.2.5 Optimizing objectives

Selecting TADs to fit each biological target quantified above is accomplished by fitting the parameters of FrankenTAD to the appropriate *OBJ* function on a given data set. These objective functions are computationally expensive to evaluate because we must run FrankenTAD with the test parameter set to find TAD sets, and then assess the objective on those TADs. We therefore use Bayesian optimization (BO) [18] to more efficiently find the parameters of Equation 1. At each step of the BO procedure, a candidate set of parameters is proposed and used to find TADs on all odd chromosomes with Equation 12. The given *OBJ* function is then evaluated on these TADs and used to inform the next candidate parameter set. We run the BO for 1000 steps, and use the parameters that produced the highest evaluation score. Because the odd chromosomes are used for parameter selection, all results are reported only on even chromosomes.

### 2.3 Data

Parameters optimizing each relevant objective function were used to predict TADs on all even autosomal chromosomes of 12 different cell types from a variety of studies (see Supplementary Table 1 for data sources) at 40kb resolution. The testing data was chosen to represent a range of biological conditions (e.g. cancerous and healthy), cell and tissue types, sequencing depths, and availability of relevant ChIP-seq data. All Hi-C data was uniformly processed from sequence data to ICE-normalized Hi-C matrices using HiC-Pro [23] (accession numbers available in Table 1). All ChIP-seq data was processed into peak formats (accession numbers available in Table 2).

**Table 1:** Hi-C data used to generate all results in this work.

| Cell type | Description | Replicates | Accession(s) | Citation |
| --- | --- | --- | --- | --- |
| IMR90 | lung fibroblast | 2 | SRR1658672, SRR1658673, SRR1658674, SRR1658675, SRR1658676, SRR1658677, SRR1658678 | [19] |
| GM12878 | blood lymphocyte | 2 | SRR1658570, SRR1658571, SRR1658572, SRR1658573, SRR1658574, SRR1658575, SRR1658576, SRR1658577, SRR1658578, SRR1658579, SRR1658580, SRR1658581, SRR1658582, SRR1658583, SRR1658584, SRR1658585, SRR1658586, SRR1658587, SRR1658588, SRR1658589, SRR1658590, SRR1658591, SRR1658592, SRR1658593, SRR1658594, SRR1658595, SRR1658596, SRR1658597, SRR1658598, SRR1658599, SRR1658600, SRR1658601, SRR1658602, SRR1658603 | [19] |
| K562 | chronic myeloid leukemia | 2 | SRR1658693, SRR1658694, SRR1658695, SRR1658696, SRR1658697, SRR1658698, SRR1658699, SRR1658700, SRR1658701, SRR1658702 | [19] |
| NHEK | epidermal keratinocyte | 1 | SRR1658689, SRR1658690, SRR1658691 | [19] |
| A549 | adenocarcinomic alveolar basal epithelial | 2 | ENCLB571HTP, ENCLB222WYT | [27] |
| LNCaP-FGC | prostate carcinoma epithelial-like | 2 | ENCLB191OGC, ENCLB473XWD | [27] |
| T47D | ductal carcinoma | 2 | ENCLB758KFU, ENCLB183QHG | [27] |
| hESC | human embryonic stem cell | 2 | SRX116344, SRX128221 | [8] |
| pancreas | tissue from 2 donors | 2 | SRX2179254, SRX2179255, SRX2179256, SRX2179257 | [21] |
| spleen | tissue from 2 donors | 2 | SRX2179264, SRX2179263 | [21] |
| skeletal muscle | gastrocnemius medialis tissue, 4 donors | 4 | ENCLB925XYW, ENCLB361HQM, ENCLB966EDS, ENCLB645GUM | [27] |
| HFFc6 | subclone of HFF-hTERT | 2 | 4DNES2R6PUEK | [6] |

**Table 2:** Accessions for all ChIP-seq data used in this work.

| Cell/tissue type | CTCF | RAD21 | H3K36me3 |
| --- | --- | --- | --- |
| IMR90 | GSM935404 | GSM935624 | GSM1055820 |
| GM12878 | GSM935611 | GSM935332 | ENCFF479XLN |
| K562 | GSM1817654 | GSM935319 | ENCFF676RWX |
| NHEK | GSM822271 | – | ENCFF253CMF |
| A549 | ENCFF543VGD | ENCFF958VNQ | GSM1003494 |
| LNCaP-FGC | GSM2827202 | – | GSM875814 |
| T47D | ENCFF903ZMF | GSM3415550 | GSM1541451 |
| hESC | GSM822297 | GSM935379 | GSM450268 |
| pancreas | GSM2827540 | – | ENCFF928PAZ |
| spleen | ENCFF459AHK | – | GSM2700497 |
| skeletal muscle | GSM733762 | – | GSM733717 |
| HFFc6 | GSM1022644 | – | GSM817238 |

## 3 Results

Each biologically-guided parameter choice was used to run FrankenTAD on each cell type for which the necessary data was available. We use JI, a metric previously used for TAD comparisons [11, 20] to quantify similarity between the TAD sets within each cell type and study the variability of TAD sets under various objectives. One cell type (NHEK) did not have replicate data, so we were unable to optimize it for reproducibility, and six of the cell types did not have publicly available RAD21 ChIP-seq data, so they could not be optimized for cohesin occupancy. We find that the JI values show a very wide distribution (Figure 2) with many high values indicating significant agreement between TAD sets, and another cluster of very low values, indicating significantly different TAD sets.

**Figure 2:**
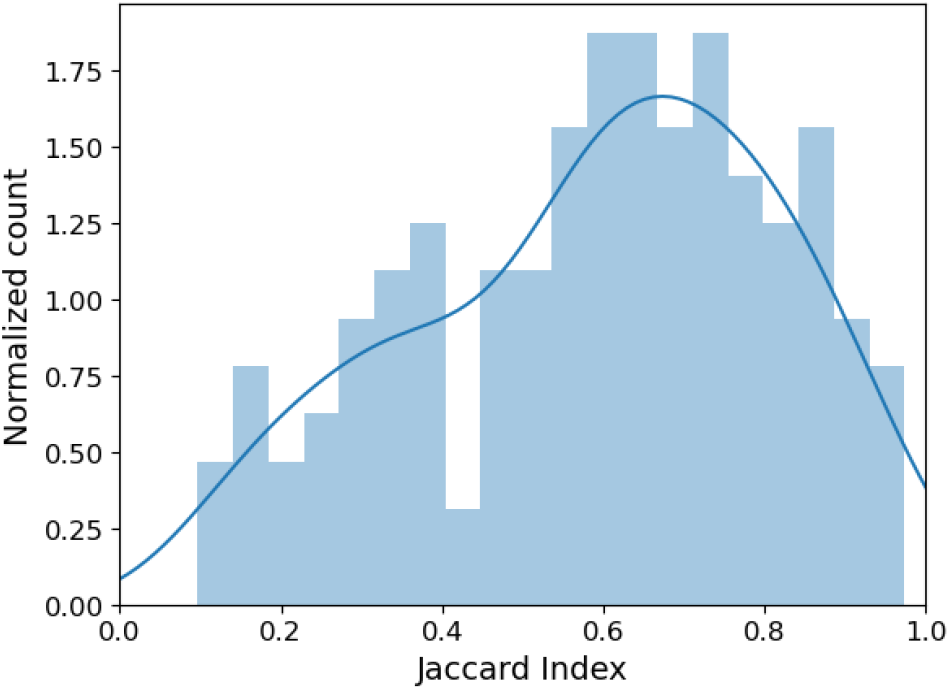
Normalized histogram of JI values across all 12 cell types and 6 objective functions.

### 3.1 Variability within cell types

The level of variability we observe in TAD sets is highly dependent on the cell type: while several showed consistency, others displayed high levels of variation. Four samples in particular, A549, HFF-c6, NHEK, and skeletal muscle tissue, showed little difference across their TAD sets (Figure 3). In each of these cell types, only one JI value lies below 0.5: between the TADs optimized for a large inter- versus intra-TAD difference, and either the TADs optimized for CTCF binding sites (A549, HFF-c6, and NHEK) or for the CTCF ChIP-seq data (skeletal muscle). On the other hand, several samples resulted in much lower JI values across their TAD sets, with IMR90 and LNCaP averaging below 0.5, and hESC and spleen tissue averaging at or below 0.55 (Figure 4). It is unclear why some samples’ TADs are highly consistent across different biological targets and others vary considerably: it does not seem to be a result of data quality or disease state.

**Figure 3:**
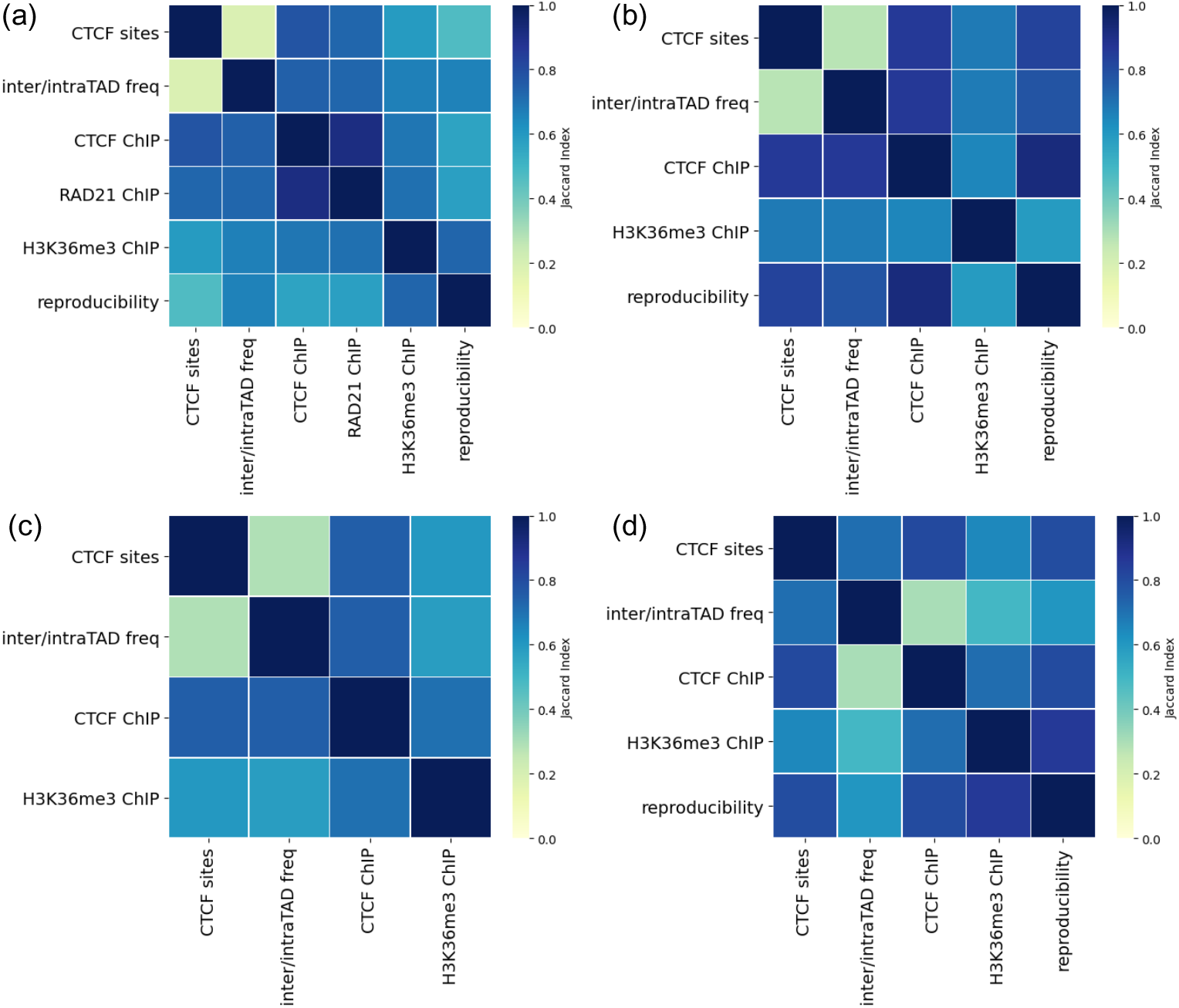
Low TAD set variability for various objective functions. Cell types: **(a)** A549 **(b)** NHEK **(c)** HFF-c6 **(d)** Skeletal muscle tissue.

**Figure 4:**
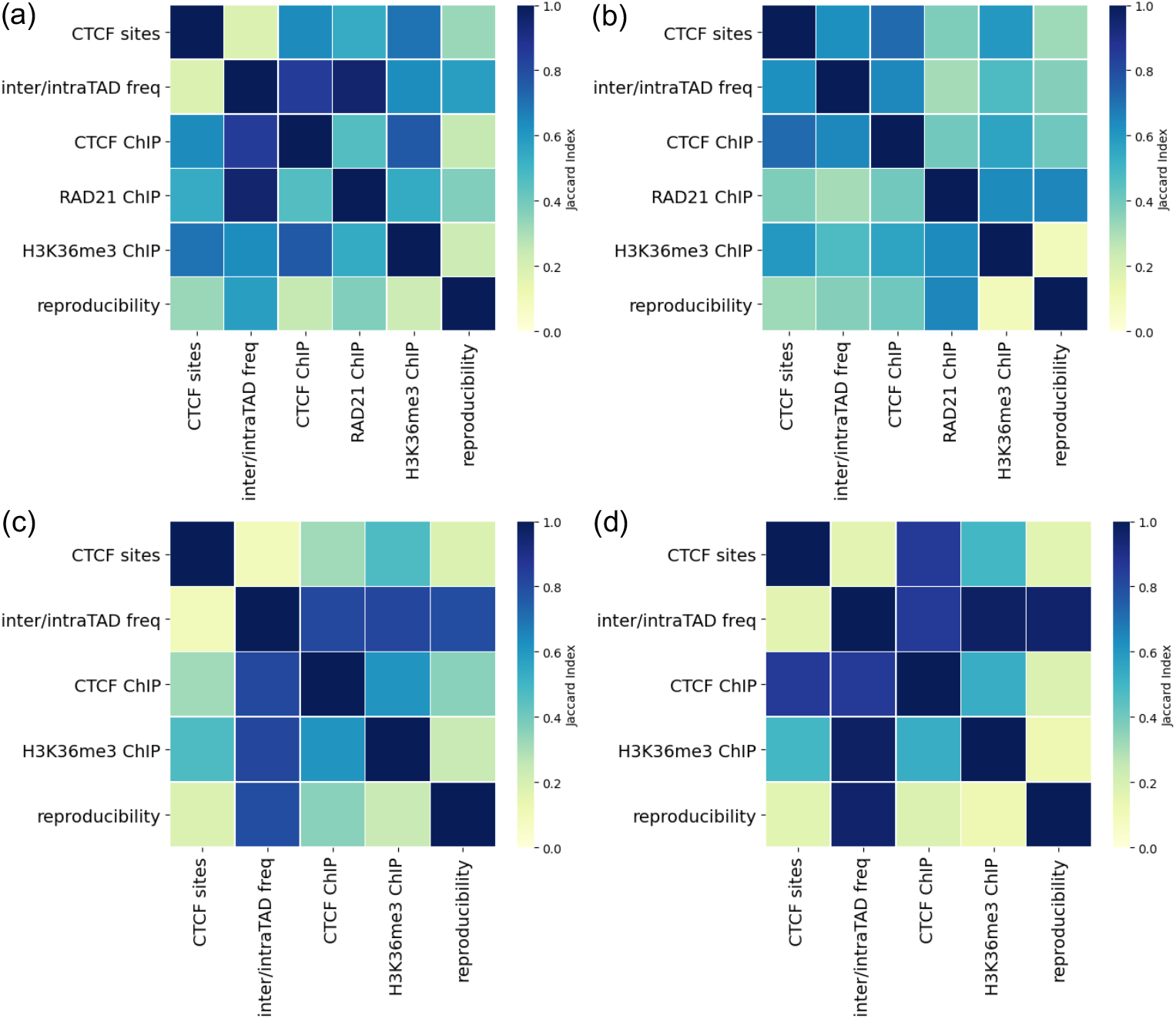
High TAD set variability for various objective functions. Cell types: **(a)** hESC **(b)** IMR90 **(c)** LNCaP-FGC **(d)** Spleen tissue.

### 3.2 Different biological targets are optimized by different sizes of TADs

One possible explanation for the variability we see in TAD sets that are optimized for different biological targets is that they represent different classes of TADs, rather than incompatible sets. It has been suggested that TADs and subTADs should be categorized more finely and defined by their mechanistic or biological components [2]. It is notable that the TADs identified for each biological target display very different distributions of TAD lengths (Figure 5). Generally speaking, optimizing for reproducibility and high inter- vs intra-TAD difference results in very small TADs, while those with high H3K36me3 at their boundaries tend to be the largest. All four protein targets lead to larger TADs, possibly suggesting that boundaries rich in protein binding are more spread out than boundaries of highly reproducible, dense subTADs.

**Figure 5:**
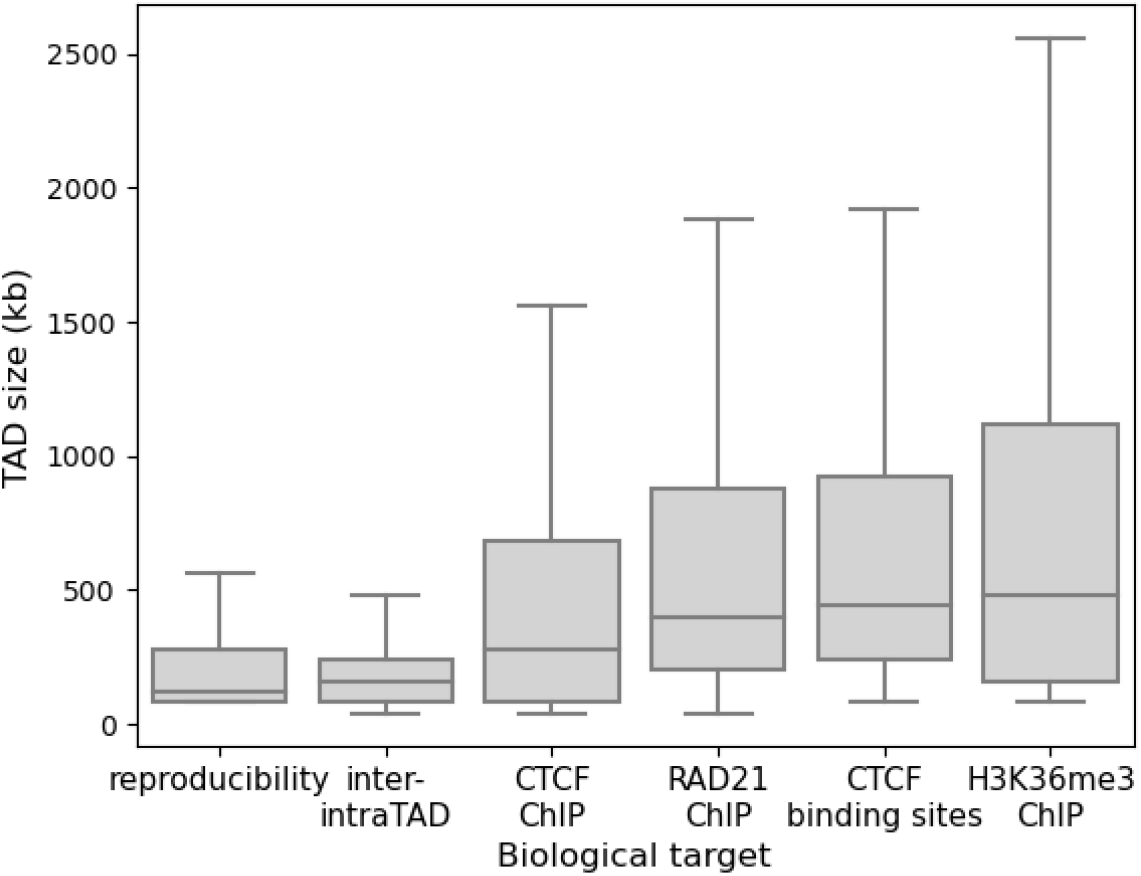
TADs optimized for different biological targets can be very different sizes.

### 3.3 Relationships between protein targets

TADs optimized for high peaks of CTCF and H3K36me3 are generally very similar to each other, but the relationship is muddled for RAD21, the cohesin subunit protein. For all but two cell types (T47D and LNCaP-FGC), the CTCF binding sites appear to be a good proxy for CTCF ChIP-seq data: TAD sets optimized for each tend to be very similar (Figure 6). Similarly, choosing parameters with CTCF ChIP-seq and H3K36me3 ChIP-seq result is generally similar TAD sets, with the exception of the pancreas tissue. Though CTCF and cohesin are believed to work together in TAD formation [13], the relationship between TADs identified by CTCF ChIP-seq and RAD21 ChIP-seq is much less clear. While two cell types (A549 and T47D) show very high similarity between TAD sets, the other four cell types have similarity values below 0.6, indicating less than 60% of TADs in common across these sets. One possible explanation comes from the loop extrusion model, which suggests that cohesin binds to DNA to create TADs, but falls off when it hits barrier elements, believed to be CTCF [12]. Under this hypothesis, we would not expect CTCF and cohesin to be frequently bound at the same genomic locations.

**Figure 6:**
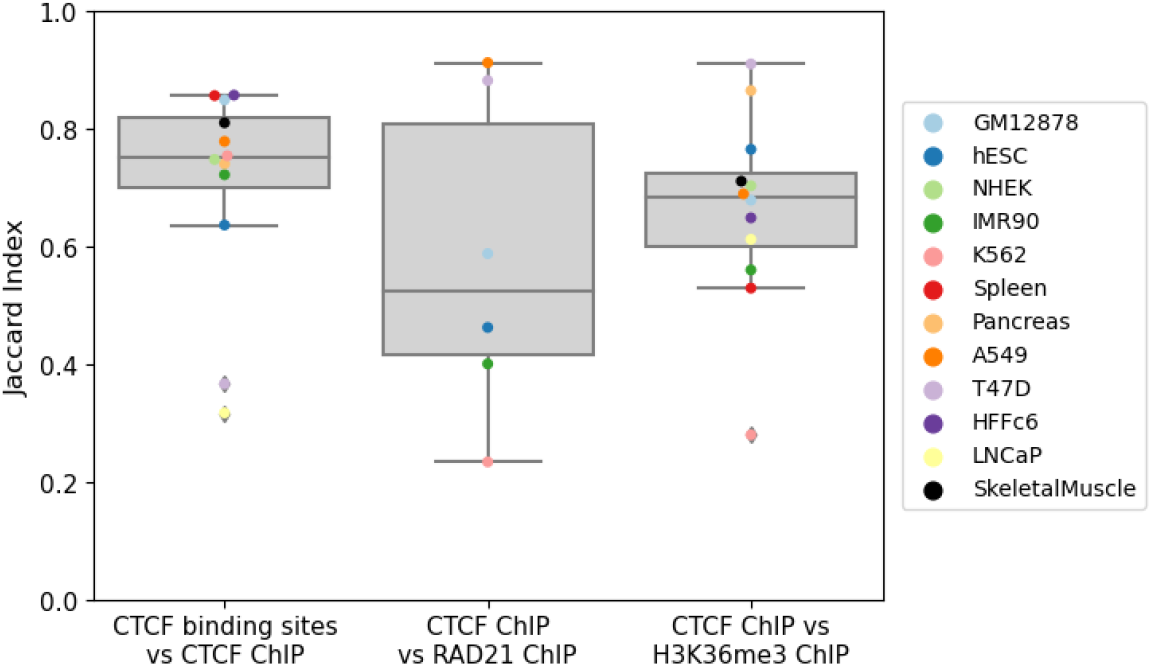
Variability of TAD sets identified with ChIP-seq or binding site data.

### 3.4 High variability in TADs from some biological targets

Looking at the variation across cell types of each objective function, we often see more variability than within a single cell type. Comparing TAD sets obtained by optimizing parameters for reproducibility, for example, shows completely different distributions in similarity values for each cell type (Figure 7). In some cell types (GM12878, K562, HFF-c6, and skeletal muscle), TAD sets optimized for reproducibility are generally similar to those optimized for other biological properties, reflected in the higher JI value distributions. In contrast, hESC, IMR90, and spleen cells produce low JI values between TADs selected by optimizing for reproducibility and those optimized for other objectives, suggesting that sometimes these objectives are not maximized by the same or even similar TAD sets. While the JI values vary significantly between cell types, in most cases the agreement between TADs identified for high reproducibility and those selected for high inter- versus intra-TAD difference is fairly high (green points in Figure 7), suggesting that these two objectives, reproducibility and high inter- versus intra-TAD difference, are generally compatible. On the other hand, the TAD sets given by parameters optimized for high reproducibility seem fairly incompatible with those given by parameters optimized for H3K36me3 peaks, shown by the lower values of the orange points in Figure 7, perhaps suggesting these TADs belong to different categories or families of TAD types.

**Figure 7:**
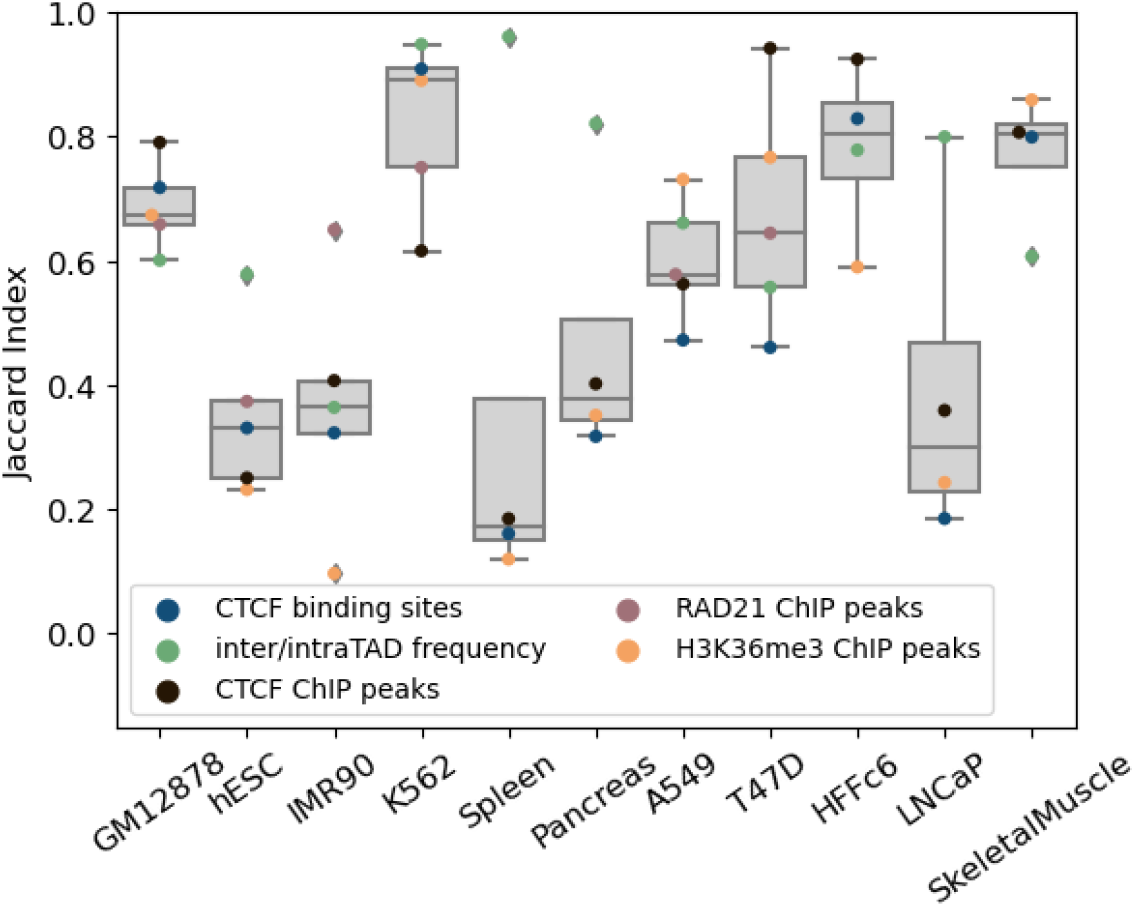
Similarity values between TAD sets optimized for reproducibility and all 5 other objective functions across cell types.

The range of similarity values between TAD sets optimized for inter- and intra-TAD frequency difference versus other objective functions also reflects significant variability. For most cell types, there was very little overlap (JI values below 0.4 for eight of the twelve cell types) in TADs produced by optimizing for CTCF binding sites and optimizing for inter- vs intra-TAD interaction differences (blue points in Figure 8). Despite these lower JI values with CTCF binding sites, TADs optimized for inter- and intra-TAD frequency difference showed much higher agreement with those optimized for CTCF ChIP-seq peaks (JI values above 0.55 for nine of twelve cell types), possibly suggesting that the binding sites are insufficient to capture both true CTCF occupancy and expected contact distribution in TAD sets.

**Figure 8:**
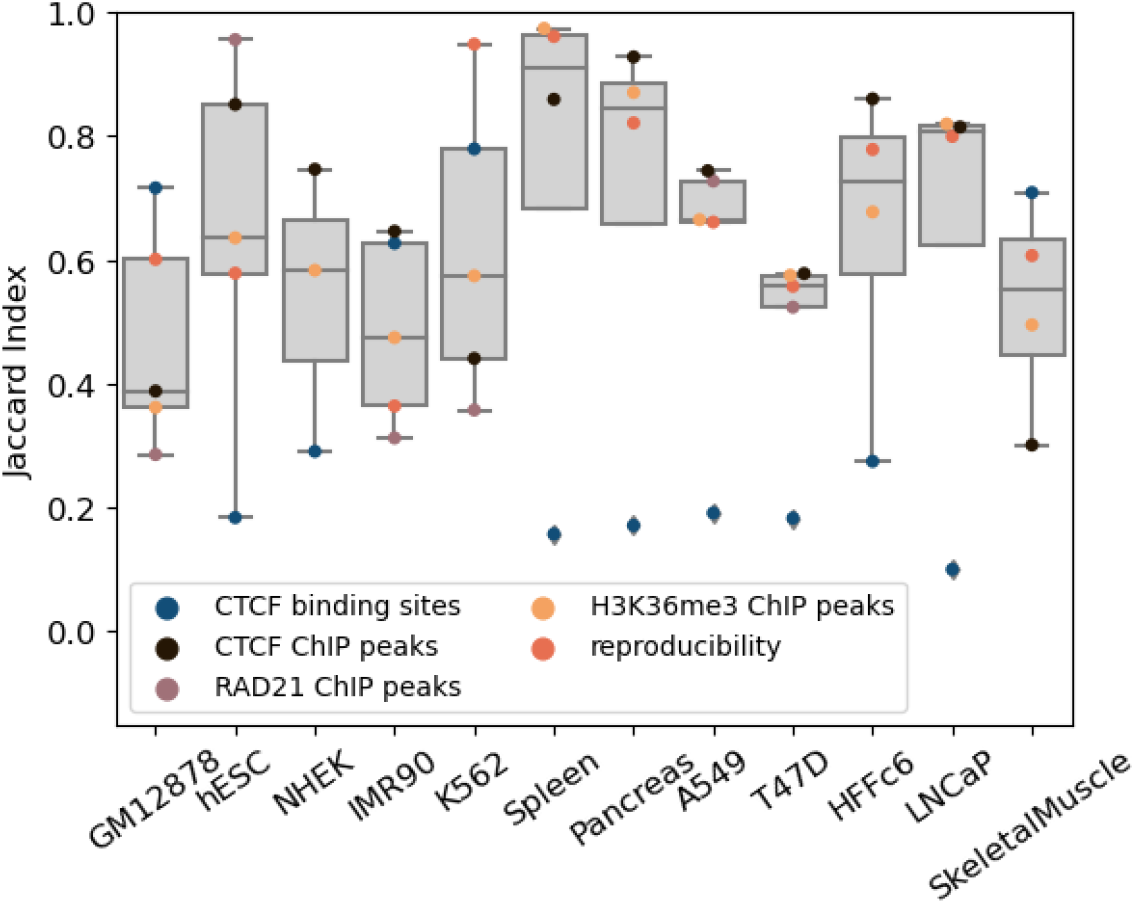
Similarity values between TAD sets optimized for high inter- and intra-TAD contact difference and all 5 other objective functions across cell types.

## 4 Discussion

We study the space of computational TAD predictions and the relationships of TAD properties with each other by developing a flexible TAD finding model with parameters tuned to a variety of data-driven properties. Our TAD-finding algorithm incorporates multiple desirable computational TAD properties, unifying many concepts for the first time under one model. Additional computational TAD definitions can be included in FrankenTAD in a straightforward manner, provided they can be efficiently computed. The algorithm design also allows for TADs to be optimized towards any quantifiable biological property through parameter tuning, therefore unifying not only computational definitions but biological patterns as well.

All epigenetic and conformational data analyzed in this work is averaged across populations of cells and therefore can only suggest trends of these populations. Our dependence on bulk data may explain some of the variability we observed. Understanding TAD-like structures in single cells is an active field of research [16, 26, 3, 1], and an analysis similar to the one presented here at the single cell level would be necessary to understand the relationships between these structures and any corresponding biological properties in individual cells. Indeed, as more single-cell Hi-C data and methods are developed, the epigenetic properties of individual TAD-like structures in single cells will likely provide significant insight into the hierarchy, sub-classes, and compatibility of TAD properties.

By choosing and comparing TADs targeted to various biological properties, we gain insight into the tradeoffs inherent in computational TAD finders. These tradeoffs seem to vary by cell type, with some cell types showing little difference in TAD sets selected for various properties while others show extreme differences. We find that optimizing the most basic TAD property, higher interaction counts within TADs than between TADs, tends to return very different, and much smaller, TADs than optimizing for any of the associated biological properties such as high CTCF or RAD21 occupancy at TAD boundaries. This suggests an inherent tradeoff between TAD properties defined solely by the Hi-C data, and those associated with other data types. These results may also suggest different families of TADs based on different properties, rather than a single TAD definition accommodating all expected biological and computational TAD properties at once.

## Financial disclosure

C.K. is a co-founder of Ocean Genomics, Inc.

## Funding

This work has been supported in part by the Gordon and Betty Moore Foundation’s Data-Driven Discovery Initiative through Grant GBMF4554 to C.K., by the US National Institutes of Health (R01GM122935), and by a Richard K. Mellon Presidential Fellowship in Life Sciences to N.S. Research reported in this publication was supported by the NIGMS of the NIH under award number P41GM103712.

